# Shadow competition and the evolution of sensory capability in sit-and-wait predators

**DOI:** 10.1101/2025.11.03.686329

**Authors:** Sumaiya Ali, Rohan S Mehta

## Abstract

Shadow competition occurs when an organism prevents another organism’s access to a resource by intercepting that resource first, thereby casting a “shadow” over the competing organism. This competition is commonly seen in the case of sit-and-wait (or ambush) predators. Shadow competition is closely related to the sensory capabilities of the organism, such as the range at which a predator can detect prey as well as the field of view of prey detection. Here, we develop an agent-based model whereby stationary, sit-and-wait predators exert shadow competition for prey on other predators on an explicit spatial landscape. Three predator characteristics can evolve: location, sensory radius, and field of view. We vary the number of available prey, the location of the prey source, the cost of sensory perception, and the energy requirements of the predator to find how shadow competition affects the evolved predator characteristics. We find that predators evolve a long, narrow sensory cone as a result of shadow competition, and that shadow competition greatly limits the efficiency of the predator-prey system. Thus, shadow competition, and the spatial interaction of predators in general, can dramatically shape predator phenotype and behavior.

## 2 Introduction

Competition, the reciprocal detriment of two individuals in the presence of each other, in its various forms is one of the fundamental concepts in ecology. Organisms compete when they both require the same resource [Birch, 1957, e.g.]. One major classification of competition is the distinction between “exploitation”, or indirect competition, and “interference”, or direct competition, but there are many conceptions of competition [Schoener, 1983]. One type of competition that crosses the line between the exploitation and interference competition is shadow competition.

Shadow competition occurs when, in the context of the physical movement of a resource or consumer, one consumer can “intercept” the resource before another consumer [Wilson, 1974, Elliott, 2002, Lubin et al., 2001, Scharf and Ruxton, 2023a]. That first consumer casts a “shadow” on the second consumer. This concept is easiest to see in predator-prey situations. Predators who ambush their prey or set traps can capture prey before another nearby predator gets a chance, depending on the location of the traps and the movement of the prey. In a river environment, predators that are upstream get the first attempt at prey compared to predators downstream. There are other examples involving situations that involve nonmotile prey or motile predators, see Scharf and Ruxton [2023a].

Few existing studies, experimental or theoretical, directly address shadow competition. In addition, those that exist consider predator locations over at most one evolutionary generation. An early theoretical study by [Wilson, 1974] proposed that antlion larvae should assume a spatial distribution of a “doughnut” shape based on considerations of group selection. The first mechanistic model, from Linton et al. [1991], directly addressed this hypothesis and outlined scenarios in which the predator population would assume a more “doughnut”-style shape or not, based on a satisficing rule in which predators would move if their current location did not meet a minimum energetic requirement. A model from Lubin et al. [2001] studied how the density and cluster size of two-lobed webs affected shadow competition; the web positions were fixed. The models of Hein et al. [2004] and Pfenning et al. [2004] address the situation where the location of the predator is restricted by the suitability of limited patches. A model from Morrell and Romey [2008] studied the trade-off between the foraging benefit of being near the periphery of a group is counterbalanced by increased predator risk at being in the periphery of the group. The recent work by Scharf [2020] builds off of the satisficing approach from Linton et al. [1991], incorporating variability in the capture radius of the predator and a correlated random walk for the prey. Notably absent in these models is the presence of evolution.

A recent review by Scharf and Ruxton [2023a] summarized the state of the field. Existing knowledge about shadow competition includes that the effect 1) increases with increasing predator density, 2) increases with decreasing straightness of prey paths, 3) should be affected by properties of the predator’s capture capabilities, and 4) should lead to the differential positioning of predators with different such characteristics among the population. In a companion paper, Scharf and Ruxton [2023b] address four modeling advances. First, they model multiple possible instantiations of the ricochet effect Rao [2009], where capture is not guaranteed and multiple encounters with predators can increase or decrease the chance of them being caught subsequently. Second, they study the effect of predator orientation. The third and fourth advances involve changes in prey behavior and are outside the scope of this study. All of the models in [Scharf and Ruxton, 2023b] focus specifically on the effect being studied, and as a consequence, the predators do not move.

In this study, we address many new threads in this field. Our main focus is to study the evolution of predator phenotypes and predator-prey interaction under shadow competition. We create an agent-based spatial model of predator-prey interaction where prey move across an arena, predators eat prey, and the predators evolve based on the results of the competition with other predators. The characteristics of the predators that evolve are position, orientation, sensory radius, and field of view. In addition, we allow predator population size to fluctuate based on their predation success, and therefore study the overall effect of shadow competition on predator population density. We find that predator sensory phenotypes consistently evolve to a relatively long and narrow sensory cone, especially when predator population density is high. We confirm previous results and expectations that predators evolve to the periphery of their habitat range, and that they generally point towards the sources of the prey. We also find that both predator population density and the efficiency of eating prey is substantially lower than what is maximally possible, demonstrating the strong effect of shadow competition in this system.

Our model is implemented in NetLogo [Wilensky, 1999] and published on NetLogo Commons at [INSERT LINK].

## Model

At the start of the simulation, each predator is randomly placed on a map of predefined size, and is assigned a random orientation (*θ*), sensory radius (*R*), and field of view (*ϕ*). All prey are added to the map at the beginning of a generation. Depending on the simulation setup, they are introduced on all four sides (4S), two opposite sides (2S; in our case left and right) or just one side (1S; in our case just left). The parameter *S* is the number of sides of prey source. Initial headings of prey are randomly chosen from any value that would not immediately take them off the map. Figure 1 shows an example diagram of two predators, their sensory areas, and four prey.

**Figure 1:**
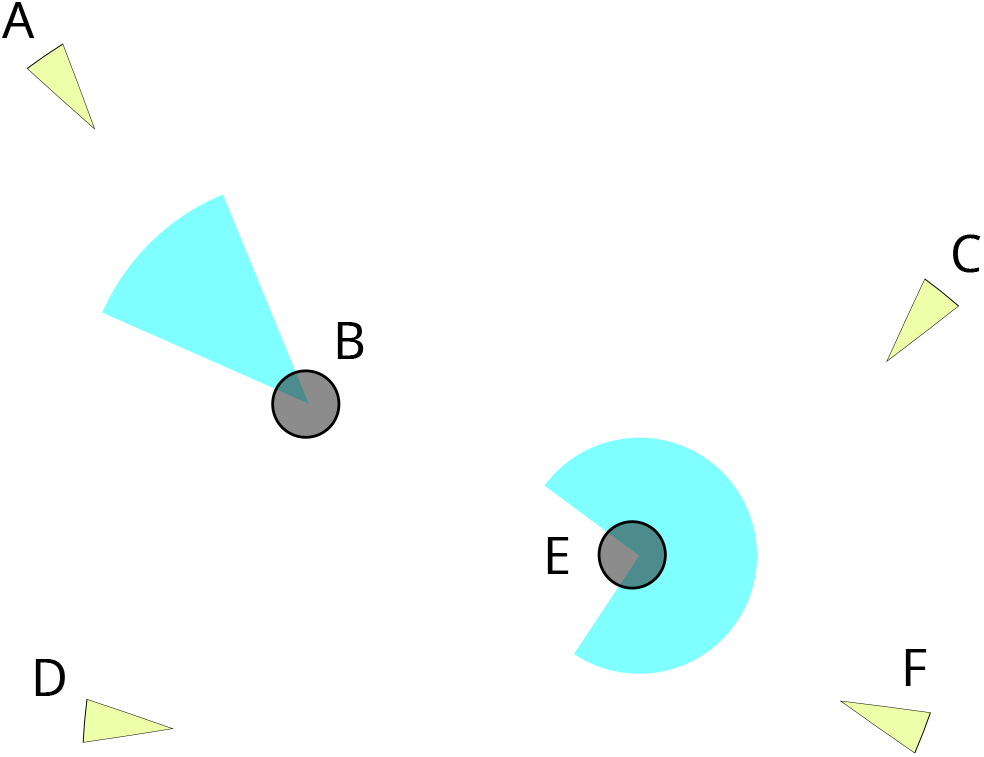
An example diagram of our model, with two predators (gray circles) at different positions and four prey (green triangles); sensory areas (aka sensory “cones”) of predators are shown in light blue. Predator B has a larger sensory radius but smaller field of view than predator E, and they have different headings. Prey A is likely to be eaten by predator B before it reaches predator E; prey C is likely to be eaten by predator E only; prey D is unlikely to be eaten by any predator, and prey F is likely to be eaten by predator E before it reaches predator B.

The NetLogo map was sized to a 41×41 square (with a tunable parameter that describes half a side length). The predators were only allowed to inhabit a smaller 31×31 square centered at (0,0) (with another tunable parameter that also describes half a side length of this smaller square). This choice allows predators to look outside their own inhabitable range. During preliminary simulations, we observed a very strong boundary effect where many predators were approaching the edge of the map and looking “inwards” because they could not “see” prey outside their habitable range, which may make sense in some biological scenarios but not in others. Hence, while we have the option of preventing predators from looking outside their range in our model, for this analysis we have allowed predators to capture prey outside of their habitable range if they can detect them.

A generation continues until all prey either walk off the map or are eaten. For a single simulation time step, the following occurs in order:

1. All prey move forward by 1 distance unit in the direction of their heading.
2. All predators detect any prey within their sensory cone.
3. Each predator, in random order, eats one random prey from any that it detected in the previous step. For each predator, this event occurs with a specified probability.

Once the generation is complete, the predators undergo a death-birth selection process. The simulation runs for *G* generations. Note that our model does allow for the ricochet effect if the probability in step 3 is less than 1; however, to keep our analyses as simple as possible, we fixed this probability at 1.

The net benefit of predator *i* is

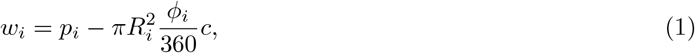

where *p*_*i*_ is the number of prey eaten by predator *i, R* is the sensory radius, *ϕ* is the field of view in degrees, and *c* is a cost constant that represents the number of prey needed to recuperate a unit cost of sensory area. For a predator with net benefit *w*_*i*_, the number of offspring it produces is

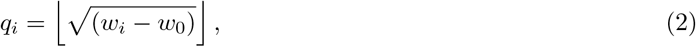

if *w*_*i*_ *>*= *w*_0_ for some net benefit *w*_0_, and 0 otherwise. The larger the value of *w*_0_, the stronger the selection. Note that *w*_0_ corresponds to the minimum amount of energy necessary to maintain growth; anything excess can be devoted to reproduction. The square root function was chosen to be sublinear but not quite as slow growing as logarithmic; preliminary simulations using a logarithmic function did not have noticeably different results.

The selection process for predators is as follows:

1. All predators with *w*_*i*_ *< w*_0_ die.
2. For each remaining predator *i*, create *q*_*i*_ new predators as offspring. These offspring mutate according to the following scheme:
  a. *R* → *R* + *N* (0, 2)
  b. *ϕ* → *ϕ* + *N* (0, 2)
  c. *θ* → (*θ* + *N* (0, 2)) mod 360
  d. *x* → *x* + *N* (0, 2)
  e. *y* → *y* + *N* (0, 2)

Note that if, after mutation, *R <* 0 then we set *R* = 0, if *ϕ <* 0 then we set *ϕ* = 0, and if *ϕ >* 360 then we set *ϕ* = 360. Also note that if the new *x* or *y* coordinate goes off the designated predator range, we set it to the edge of the that range. Finally, note that there are overlapping generations; if a predator ate exactly *w*_0_ prey, it would still survive but produce no offspring.

We performed three parameter sweeps: one where we allowed both the radius *R* and the field of view *ϕ* to mutate with a finer grid of parameters (the “free mutation” sweep), and one where we restrict *R* or *ϕ* to mutate over a coarser grid (the “fixed-radius” and “fixed field of view” sweeps). Parameter ranges for these sweeps are found in Table 1. Each simulation was initialized with 150 predators. When the sensory radius *R* was allowed to mutate, initial values of *R* were chosen from a uniform distribution between 0 and 5. When the field of view *ϕ* was allowed to mutate, initial values of *ϕ* were chosen from a uniform distribution between 0 and 360. We removed simulations where there were no predators after 100 generations: leaving us with 6638 out of an original 9000 fixed field of view simulations, 4073 out of 5400 fixed-radius simulations, and 24813 out of 30000 free mutation simulations.

**Table 1:**
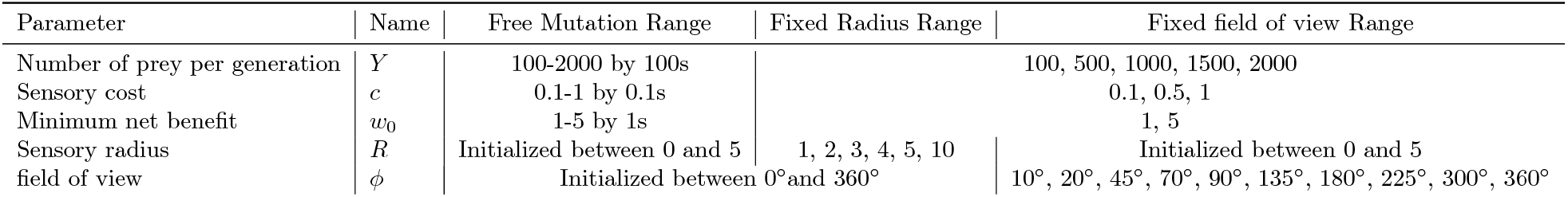
Parameter ranges used in simulations for parameters whose ranges varied in the simulations. All options for the number of sides of prey sources *S* (1, 2, and 4) were used for all sweeps.

## Results

### General Effects of Model Parameters

Table 2 describes ANOVA results for percent variance explained of the four main model parameters (prey per generation *Y*, sensory cost *c*, minimum net benefit *w*_0_, and the number of sides of prey source *S*) and various measurements obtained from each simulation. When applicable, Table 2 also displays the direction of the effect of the parameter (“nm” means “non-monotonic”, “+” means “monotone increasing”, and “-” means “monotone decreasing”; note that these are not exact, but general descriptions of the trends). For reference, a summary plot of the means of these measurements compared to the model parameters is provided in Figure 2 in order to visualize the trends. All p-values of these ANOVA results were less than 1 *×* 10^−26^; this is to be expected in a simulation study set up in this way [White et al., 2014], and instead we focus on percent variance explained as a rough measure of effect size. The specific measurements in Table 2 and Figure 2 are discussed in subsequent sections.

**Table 2:** ANOVA results for testing the effects of the four main model parameters on various different measurements obtained from simulations. Each measurement is obtained by average over all individuals in a particular simulation. All p-values are less than 1 *×* 10^−26^. The columns after each percent variance explained refer to the direction of the effect: “nm” means “non-monotonic”, “+” means “monotone increasing”, and “-” means “monotone decreasing”. MDFC = Mean Distance From Center (of the map). Effect directions are visualized in Figure 2.

| Parameter | Radius | Field of View | Sensory Area | Number of Predators | Mean Prey Eaten | Total Prey Eaten | MDFC |
| --- | --- | --- | --- | --- | --- | --- | --- |
| $Y$ | 0.03 nm | 0.07 nm | 0.07 − | 0.38 + | 0.03 nm | 0.69 + | 0.01 nm |
| $c$ | 0.00 − | 0.05 − | 0.34 − | 0.02 − | 0.01 + | 0.02 − | 0.04 − |
| $w_0$ | 0.06 + | 0.03 + | 0.08 + | 0.35 − | 0.72 + | 0.10 − | 0.33 − |
| $S$ | 0.04 + | 0.00 nm | 0.06 + | 0.02 − | 0.05 + | 0.00 + | 0.10 − |
| Residuals | 0.87 | 0.85 | 0.46 | 0.24 | 0.20 | 0.19 | 0.52 |

**Figure 2:**
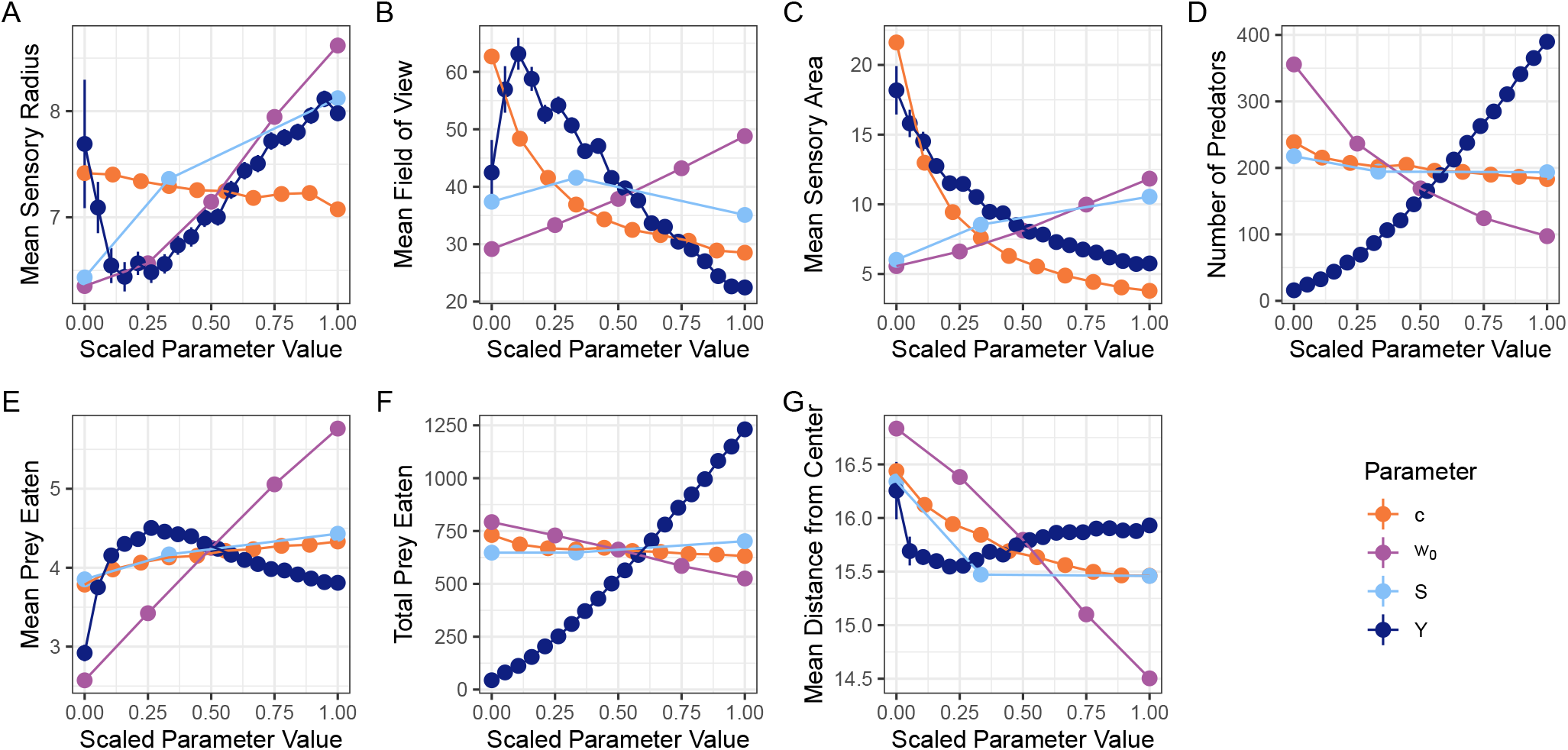
The effects of various parameters on various measurements corresponding to those provided in Table 2. The x-axis is the value of the parameter scaled so that 0 is the minimum value of the parameter and 1 is the maximum value of the parameter. Error bars correspond to standard errors. Points are averages over measurements taken by averaging (or summing, in the case of total prey eaten) over each individual in a simulation.

### Spatial Distribution of Predators

The distribution of predators in space is in general towards the boundary of the map and the sources of the prey, but the extent to which this is the case depends on the parameters. The predators are generally oriented towards the source of the prey.

Figure 3 displays heatmaps of the x-y coordinates of all predators in all simulations from the mutation regime comparison sweep. The predators are concentrated at the sources of the prey, but occasionally there are predators more towards the center of the map as well.

**Figure 3:**
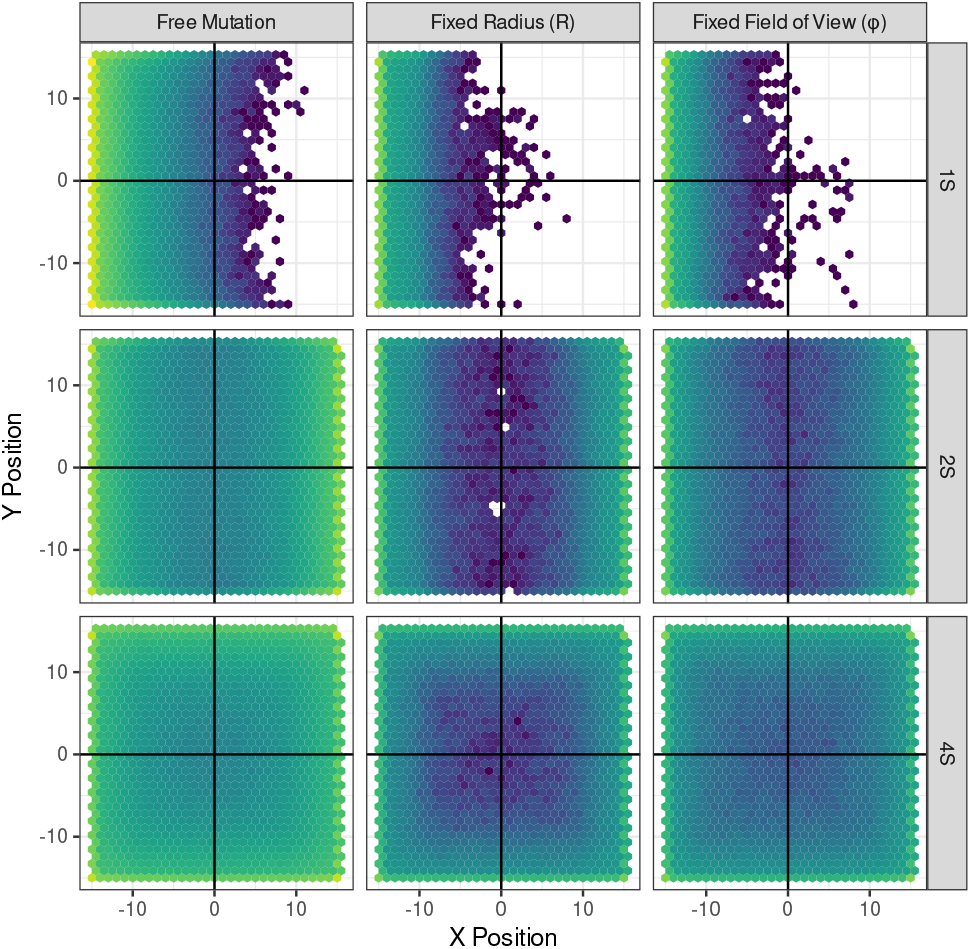
Heatmaps of all the x-y positions of the predators in all simulations. The vast majority of predator population size is towards the edge of their range and towards the source of the prey. Note that the color scale here is log-transformed and that the intensity of the color is lower for the simulations with fixed mutation because there were fewer parameter sets in that regime; the overall pattern is the same in all regimes.

From Table 2 and Figure 2G, there is one main determiner of how far, on average, the predator is from the center of the map (MDFC, or Mean Distance From Center (of the map)): as *w*_0_ increases, predators tend to move closer to the center and away from the edges. The same effect is seen for *c* and *S*, but to a smaller extent and in a saturating manner. For *Y*, it appears that for extremely small numbers of prey, predators are closer to the edge, but as the number of prey increases, predators move away from the center and then slowly move a little bit back towards the edge. Note that all of these effects are relatively small; even the biggest range here is about 2.5 distance units in a 31×31 grid, and the vast majority of predators are near the sources of the prey.

The distribution of headings for all simulations in aggregrate, separated by number of sides and which variables are allowed to mutate, are shown in Figure 4. For 1S, predators point directly at prey sources. For 2S, predators point mostly towards prey sources, but for 4S something interesting happens: for fixed *ϕ*, predators point everywhere with no bias. For fixed *R*, predators generally point towards the prey sources. But for free mutation, predators point to the corners. As it turns out, there is a large shift in predator pointing behavior for larger values of *R* and small values of *ϕ* (see Supplementary Figures **??** and **??**). When *R* is fixed, for small *R*, predators point to the sides. But for larger *R*, they point to the corners. When *ϕ* is fixed, for large *ϕ*, there is no bias in heading direction. For small *ϕ*, predators again point to the corners. This change is probably due to the fact that a large *R* with a narrow *ϕ* a certain distance away from the edge of the map pointing to a corner will have more “space” to work with than pointing towards the edge, as the map is a 2D square. Indeed, as we shall see in Figure 7, the evolved values of *R* are large enough that we would expect to see this corner-pointing behavior.

**Figure 4:**
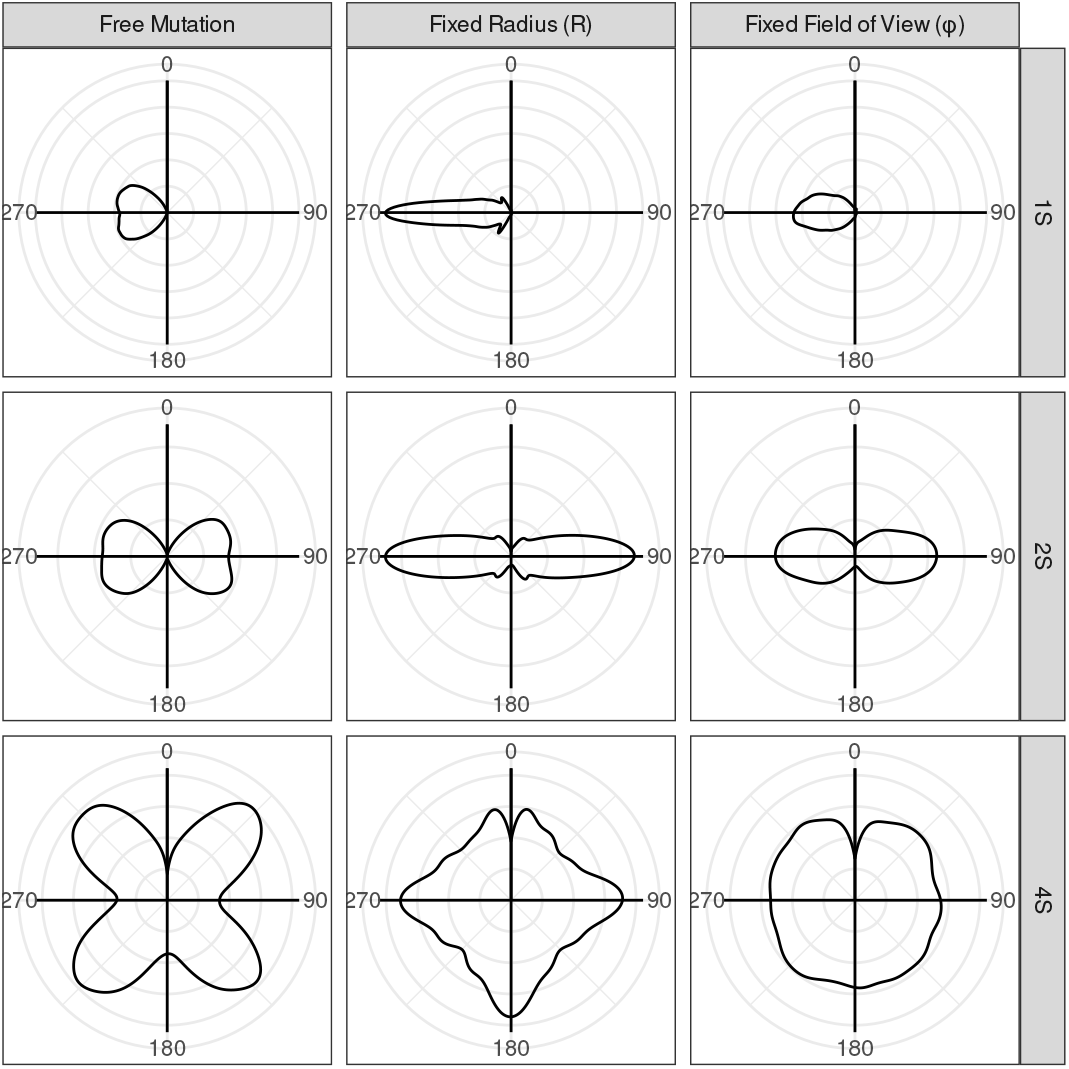
Predators generally point directly to their prey source (to the left for 1S, to the left and right for 2S, and in all four cardinal directions for 4S), or to some average of the prey sources (for free mutation and 4S, the predators point to the corners). The “pinch” points at 0 in the last two panels are artifacts of the kernel density estimation.

### Predator Population Size

For this section, we focus on the situation where both sensory radius *R* and field of view *ϕ* are allowed to mutate.

From Table 2 and Figure 2D, the number of predators is dramatically affected by the number of prey per generation *Y* (positively) and the minimum net benefit *w*_0_ (negatively), with comparatively little effect of sensory cost *c* or the number of sides *S*.

When considering predator population size, it is useful to think about the maximum possible predator population size.

For any particular simulation, the maximum number of prey that can be eaten per generation is the total number of prey entering the map *Y*. Because prey are initialized outside the predator range, some of them can “miss” the predators entirely (although they may be able to be caught by predators on the edge of their range). Thus, in practice, the “best” that these predators can do is usually less than capturing all *Y* possible prey. Appendix A computes this value for our map setup, which is approximately 74%. So we consider *Y* ^*′*^ = 0.74*Y* for following analyses. Note that this change only affects the scaling in Figure 5; all other results are basically the same as if we had just used *Y*.

**Figure 5:**
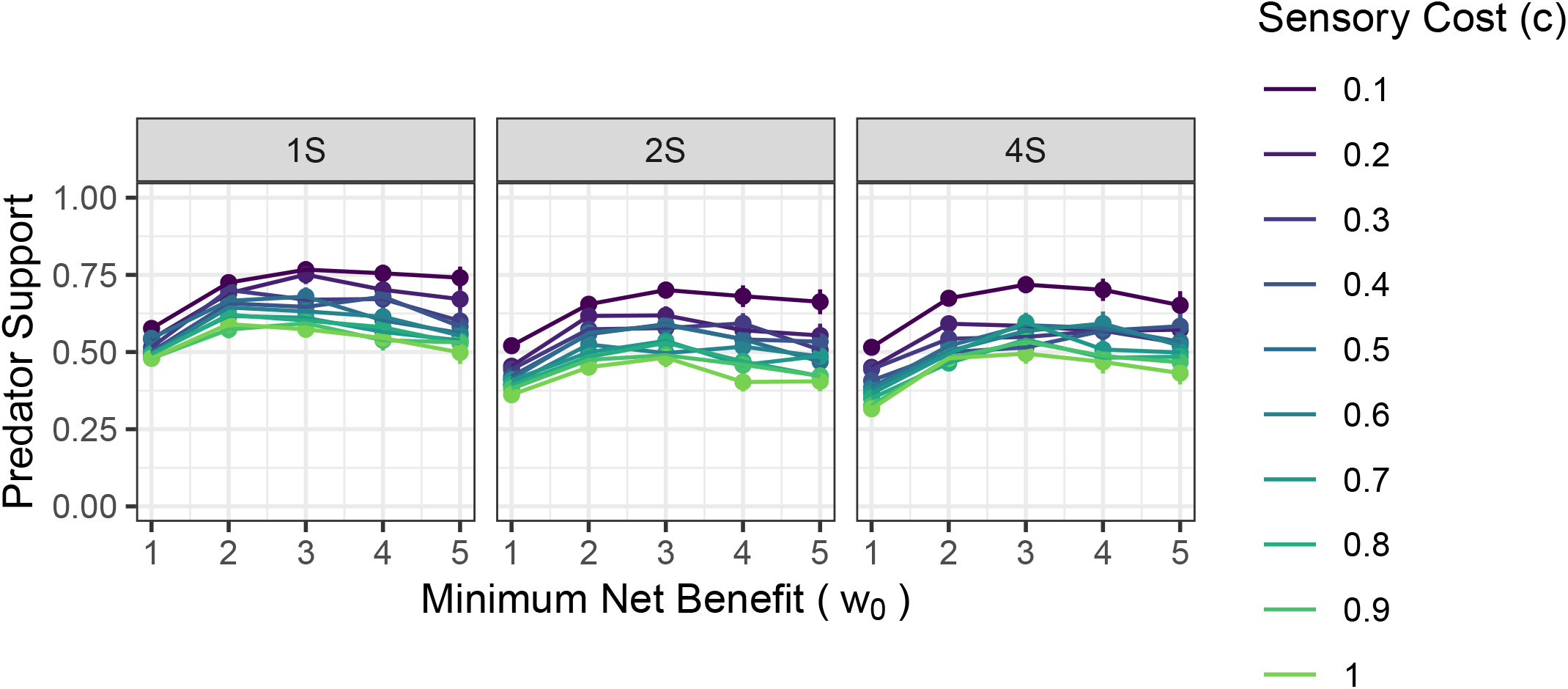
The “predator support” (the limit of the ratio between the number of predators and the maximum number of predators for a given set of parameters as the number of prey *Y* ^*′*^ → ∞) as affected by sensory cost *c*, minimum number of prey eaten *w*_0_, and number of sides of the prey source. Each point is a slope estimate from a linear regression of *P* onto *Y* −*w*_0_, multiple by *w*_0_. Error bars are *±*2SE. The largest effects are that of cost (which is consistent), between *w*_0_ = 1 and *w*_0_ *>* 1, and between 1S vs 2S and 4S.

The maximum possible predator population size is is obtained by giving each predator the minimum amount of prey possible *w*_0_:

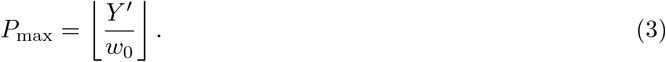

Predator population size *P* is approximately linear in the number of prey per generation *Y* (see Figure 2D and Supplemental Figure 10A), especially for large *Y*. We can therefore model P as

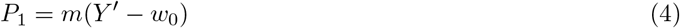

For some slope *m* that depends on the other parameters of the system. Note that for a predator to exist, there has to be at least *Y* ^*′*^ *> w*_0_, hence the shifted model. The ratio of *P*_1_ to *P*_max_ tells us how much the parameters support a predator population given a particular number of prey per generation; a lower value would indicate increased competition or increased sensory cost:

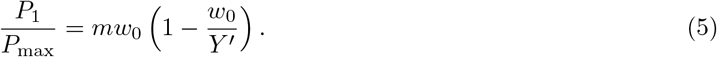

Eq. 5 predicts that as *Y* ^*′*^ → ∞, the ratio 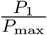 will saturate to some value *mw*_0_, where *m* is from the linear relationship of *P* to *Y* ^*′*^ (eq. 4 and Supplementary Figure 10A and B). Indeed, this is what we observe (Supplementary Figure 10C). We can infer this slope *m* by linear regression. We call this slope the “Predator Support”, and it indicates how much of the maximum number of predators a system can support when there are a large number of prey available. This value is shown in Figure 5.

In general, increasing cost decreases predator support. Increasing *w*_0_ actually increases predator support initially (by potentially alleviating competition between predators, shifting the pressure to satisfying metabolic needs), but too high a *w*_0_ and you set too high a bar for predators. The one-sided prey situation has higher predator support than the other prey source situations, suggesting that there is more interference between predators when prey come from all sides. The range of predator support goes from about 30% of the maximum to about 75% of the maximum. There is therefore a substantial effect of interpredator competition even at low sensory cost and a restrictive survival criterion (high minimum net benefit).

### Number of Prey Eaten

Table 2 and Figures 2E and F depict two different measures of the number of prey eaten: the mean prey eaten per predator, and the total prey eaten, respectively. The mean prey eaten per predator is only dramatically affected by *w*_0_, which makes sense because that *w*_0_ sets a lower bound on how much a predator can eat to survive. The total prey eaten, on the other hand, depends extremely strongly on mostly the prey per generation *Y*.

As with the number of predators, we can try to rescale the number of prey eaten to gain more insight into these effects.

Assuming that every existing predator can survive to the next generation (which is not in general true), the minimum total number of prey eaten in a particular generation for a specific number of predators *P* is *w*_0_*P*. The maximum number of prey eaten is *Y*, and with the missing correction *Y* ^*′*^. We can thus rescale the total number of prey eaten to be in between these values by a linear combination of the maximum and minimum: let *p*^*′*^ be the scaled value of the total number of prey eaten:

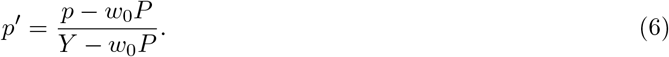

Figure 6 plots several different quantities relevant to the quantity *p*^*′*^. Figure 6A shows the whole distribution of *p*^*′*^ over all free-mutation simulations; this distribution is very narrow and rarely ventures above 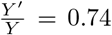 (the dashed vertical line) or below 0. Values above 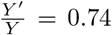 suggest that the predators are successful at “reaching over the barrier” and grabbing some prey outside their habitable range, and values below 0 indicate that some predators in the final simulation generation did not eat enough prey to survive to the next generation. Because the distribution of *p*^*′*^ is not strongly affected by any parameters other than *w*_0_, Figures 6B-D show instead the fraction of simulations where *p*^*′*^ *<* 0 for *c, Y*, and *S*, where there is an effect: increasing *Y*, decreasing *c*, and decreasing *S* all increase this fraction, which makes sense as the first two situations would lead to more predators and therefore more competition, and the third situation has only one prey source and therefore more competition.

**Figure 6:**
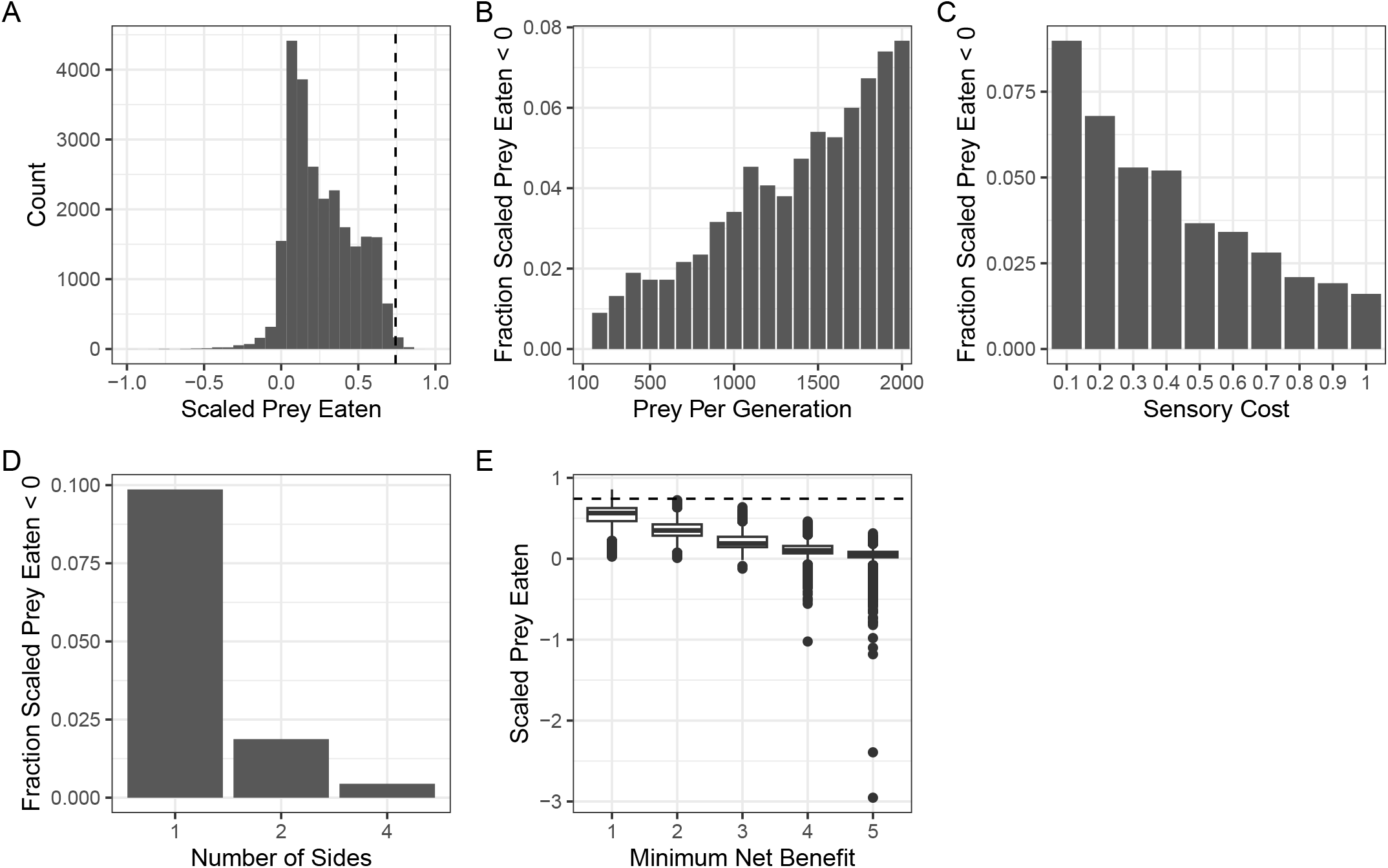
Effects of model parameters on scaled prey eaten *p*^*′*^ (eq. 6). (A) The distribution of scaled prey eaten over all free-mutation simulations. 1 means that all *Y* prey were eaten. The vertical dashed line is at 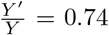. 0 indicates *w*_0_*P* prey were eaten. Negative numbers indicate that some predators ate fewer than *w*_0_ prey. (There are two simulations that go out to −2 and −3 on this plot but the axis was cut off for readability. They appear in E.) B-D) The fraction of simulations with *p*^*′*^ *<* 0 for varying (B) *Y*, (C) *c*, and (D) *S*. (E) The effect of *w*_0_ on the distribution of *p*^*′*^. The horizontal dashed line is at 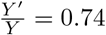.

Figures 6E shows the only parameter effect on *p*^*′*^ that notably shifts the whole distribution: *w*_0_. When *w*_0_ = 1, *p*^*′*^ is close to 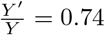 (the dashed horizontal line). When *w*_0_ = 5, *p*^*′*^ is very close to its ostensible minimum (0, which in this case means that all predators eat close to *w*_0_ prey) and there is a notable increase in the tail of the distribution where many predators will not survive to the next generation.

### Phenotype Evolution

Figure 7 depicts the *R* − *ϕ* phenotype space along with evolved phenotypes for simulations with free *R*, free *ϕ*, or both. In general, the evolved phenotypes for the non-free-mutation scenarios fall along a curve where the evolved areas are relatively similar (reflected in Figure 7 with the curve that represents the average area across all restricted simulations) and the free mutation scenarios are also dispersed along this curve, though there are higher concentrations at different parts. Importantly, the highest concentration of results is at a smaller-than-10-degree *ϕ* and a radius between 5 and 10, indicating a general tendency for narrow, long sensory cones.

**Figure 7:**
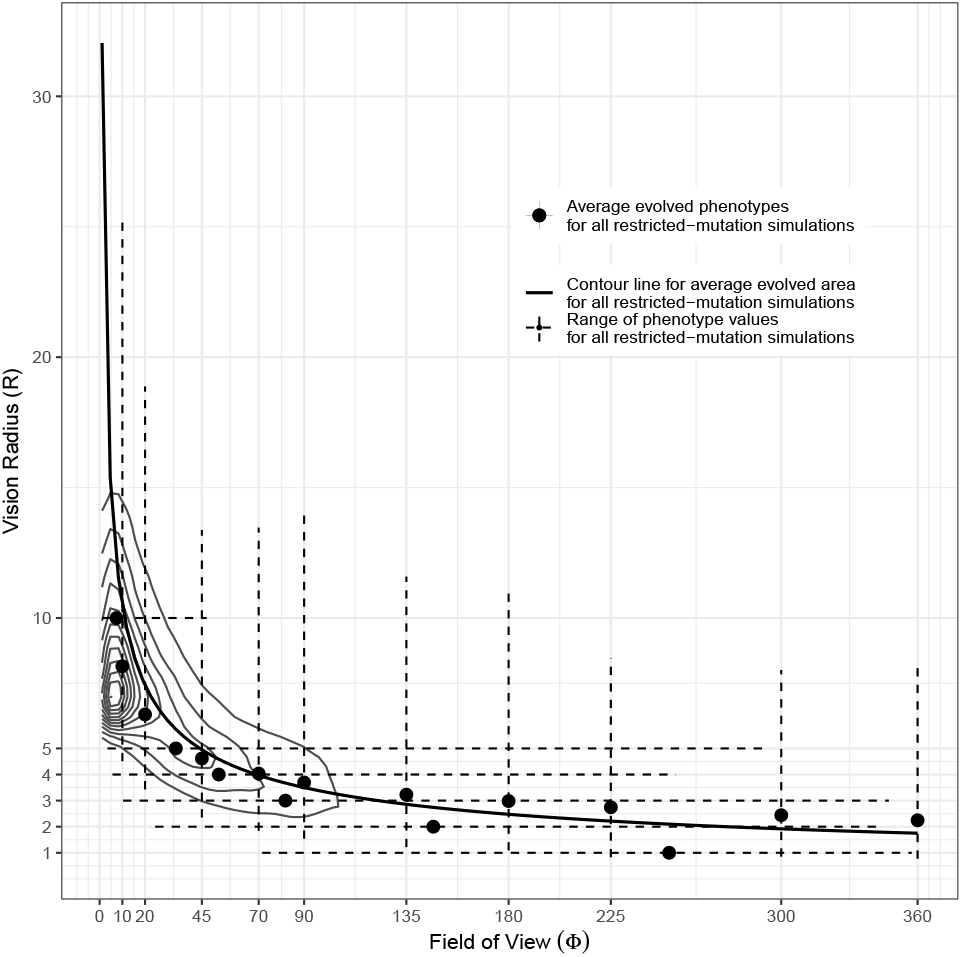
Evolved phenotypes across free-mutation and restricted-mutation simulations in the mixed-mutation regime sweep. The transparent dots are evolved phenotypes for all simulations where both radius *R* and field of view *ϕ* were allowed to mutate; averaged over each simulation. The solid dots are evolved phenotypes for all simulations where either *R* or *ϕ* was not allowed to mutate, averaged over all simulations. The dashed lines are the range of values of the other phenotype variable that these restricted-mutation simulations reached (*R* for fixed *ϕ*, and *ϕ* for fixed *R*); the minima and maxima were computed over the average values from each simulation. Finally, the solid line is the value of *R* that corresponds to a particular value of *ϕ* given a fixed evolved sensory area, which is in this case the evolved area for each restricted-mutation simulation averaged over all individuals in each simulation.

From Table 2, the model parameters themselves do not have a large effect on specific values of radius or field-of-view. From Figure 2A and B, we can see that there are small directional effects of *c, w*_0_, and *S*, as well as a small nonmonotonic effect of *Y*, but these effects do not, in general, take the phenotype out of the dense part of its distribution in Figure 7. However, the model parameters do have consistent and substantial effects on the overall evolved area (Figure 2C). In general, area decreases with increasing *Y* and *c* and increases with increasing *w*_0_ and *S*.

If sensory area is the main evolutionary outcome and the radius and field-of-view separately are not, then why is the phenotype distribution so narrowly concentrated in one spot instead of along the whole curve that would represent the evolved area? More predators should lead to more shadow competition, so if we consider how these phenotypes change with the number of predators, that might offer us a clue.

Figure 8 shows the effect of the number of predators on *R* and *ϕ*. In general, when there are more predators, evolved values of *R* and *ϕ* converge, and when there are fewer predators, many values of *R* and *ϕ* can evolve. As is shown by the accompanying densities, the values at which *R* and *ϕ* converge are close to the overall modes of the distributions. This value matches that of the field of view that most of the simulations attain, but is notably higher than the value of the radius that most simulations attain.

**Figure 8:**
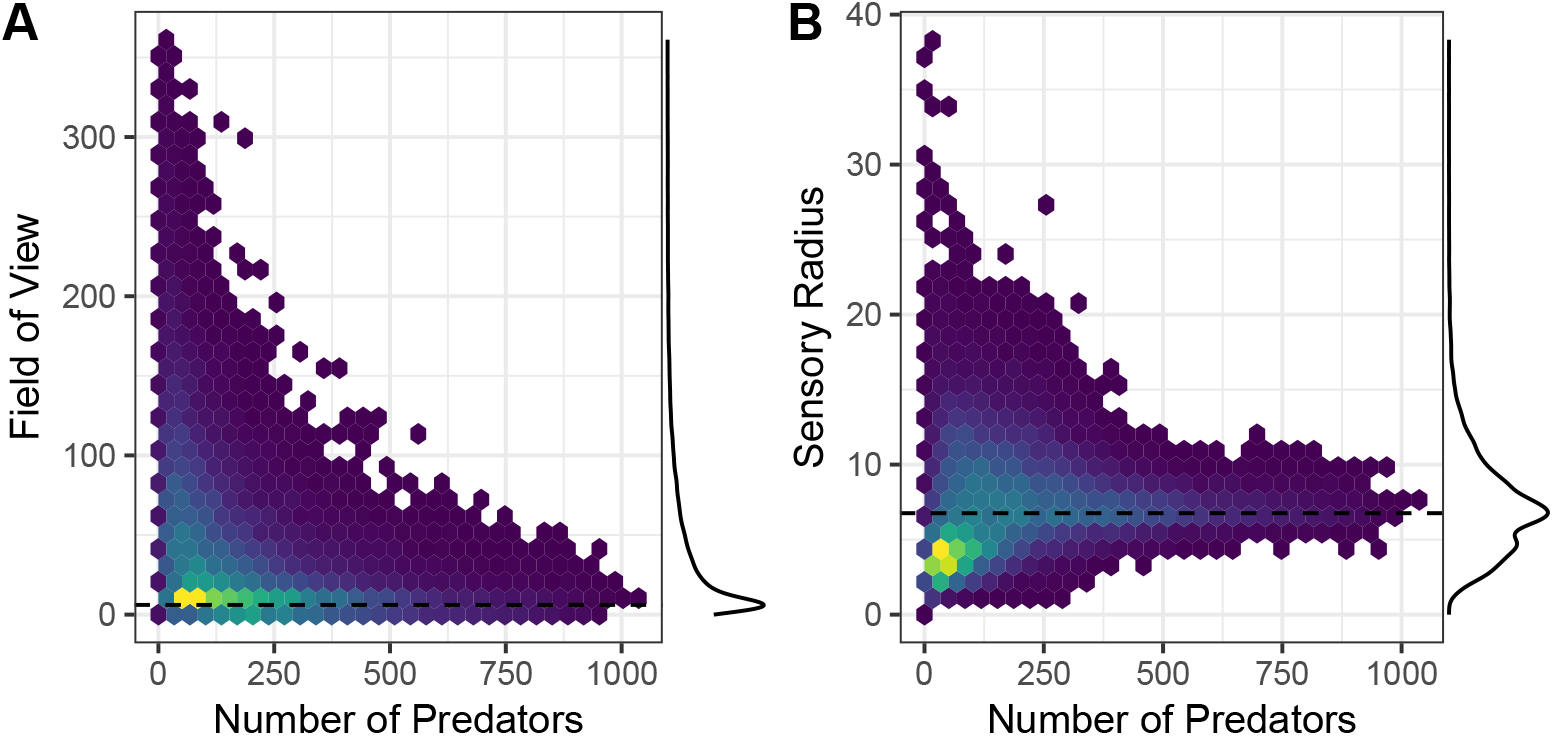
As the number of predators increases, the range of evolved and fields of view (A) and sensory radii (B) narrow. The modes of the marginal densities of field of view and senroy radius are displayed by horizontal lines. Each plot is a hexagonal grid of a scatterplot with each data point a free-mutation simulation.

## Discussion

We have developed an evolutionary agent-based model of shadow competition in a simple predator-prey system and analyzed it over a wide range of the four main model parameters: the number of prey introduced in each generation, the cost per unit area of maintaining a particular sensory system, the minimum amount of energy needed to survive, and the number of disconnected sources of prey. The heritable characteristics of the predator that could evolve were: its location in a restricted habitat range, its orientation, its radius of sensory perception, and its field of view. The number of predators in a particular generation was allowed to vary based on a death-birth process with overlapping generations where all predators that did not meet a minimum energy requirement died and all other predators produced offspring based on how many prey they ate and how much their sensory area cost.

In general, predators evolved to be close to the edge of their habitat range, pointing towards their prey sources. The extreme “doughnut” nature of this spatial distribution is potentially too extreme to be observed in nature. One aspect of our model that is included in the implementation but not tested in this study (in order to keep it somewhat simple) is the “ricochet effect”, which is where the probability of capture upon sensory perception is less than 1. One might expect the spatial distribution of predators to be less “doughnutlike” and more even when the capture probability is low. In addition, other tradeoffs may exist for individuals at the periphery of the distribution that we do not account for.

When the prey come in from all sides, predators point to the corners instead of the sides when their sensory radii are large and their fields of view are small. This effect would go away in a circular range, but it is worth noting that actual predator habitat ranges probably have “corner”-like features, and so it is worth considering how predators would react to being on the edge of their habitat.

The evolution of the phenotype itself is mostly based on the total sensory area rather than the individual fields of view and sensory radii that compose that area. This area decreases when there are more available prey (which means there are more predators, and so more competition), when there is an increased cost to producing that area, when prey come from fewer sides (less available space and therefore more competition), and when the minimum amount of energy needed to survive is low (again, more predators so more competition).

In this relatively extreme shadow competition situation, the predators settle on essentially one major phenotype strategy: a narrow field of view and a relatively large sensory radius. This result can be interpreted as the field of view evolving to be narrow due to shadow competition, and the radius evolving to increase to compensate so that sensory area is large enough to find prey.

Predator population size appears to be well below what one might expect to be its maximum given the amount of prey provided and the minimum energy requirement, suggesting a high level of competition. The population size appears to mostly depend on the number of prey available and the minimum amount of energy needed to survive, depending very little on sensory cost. However, sensory cost plays a much larger role in determining the actual phenotypes. This result suggests that the evolution of sensory radius and field of view can potentially “mitigate” the sensory cost without having too much effect on the actual predator population size, so long as there are enough prey around. In addition, the fact that the range of evolved field of view and radius values narrows dramatically with predator population size suggests that the phenotype evolution is indeed being driven by predator population size and competition between predators.

Relatively few simulated here are “efficient” with respect to the amount of total prey eaten; the number of prey eaten is in general much closer to that which would occur if each existing predator ate the minimum amount of prey needed to survive, further suggesting strong competition. The relatively few cases where the total number of prey eaten go beyond our approximate maximum given the limited predator range compared to the prey range suggests that “reaching over the wall” to grab prey is not the norm and may not be worth the increased sensory cost. The fact that there are relatively few cases where the total number of prey eaten are not enough to maintain the current predator population size suggests that evolution has mostly “finished” determining population size in most cases; running the simulations longer may remove these cases but if the situation is extreme (very high minimum energy requirement and very uneven distribution of prey eaten among predators) these cases may always occur.

In addition to studying the ricochet effect, there are many potential extensions and modifications of our model that can be studied in the future. For instance, our model uses ballistic motion for prey; a correlated random walk or perhaps even a Lévy walk may better capture prey foraging behavior. Other aspects of this system that previous work has studied that our model does not currently have include patchy predator habitats and within-generation predator position choice. Explicit tradeoffs between sensory radius and field of view could also be included. Finally, we primarily used the fixed-radius and fixed-field-of-view simulations to better explore the phenotype space; these situations can also be studied in and of themselves as they can better match the physiological contraints or behavior patterns of existing organisms (for instance, antlion larvae and web-building spiders are essentially restricted to a panoramic field of view, whereas organisms that rely on sight or who set traps in specific places may have more limited fields of view).

Overall, our model demonstrates that over many generations, shadow competition can dramatically shape predator behavior (in terms of their locations and orientation) and sensory phenotype, with very strong shadow competition leading to longer, narrower sensory cones. In addition, our model suggests that shadow competition itself can be a very strong determiner of predator population size and of prey survival.

## Acknowledgements

This work was funded by the Creative and Scholarly Endeavors (CASE) program at Elmhurst University.

## Supplementary Information

### A: The fraction of prey that are accessible to predators

The fraction of prey that are actually accessible to a predator, given that the map (with side length *m*) is bigger than the predator habitat range (with side length *s*). Due to symmetry, we cna focus on one side of prey source: Figure 9 is a diagram of this calculation for the top side. The, possible prey headings range from −*π* to 2*π* in radians.

**Figure 9:**
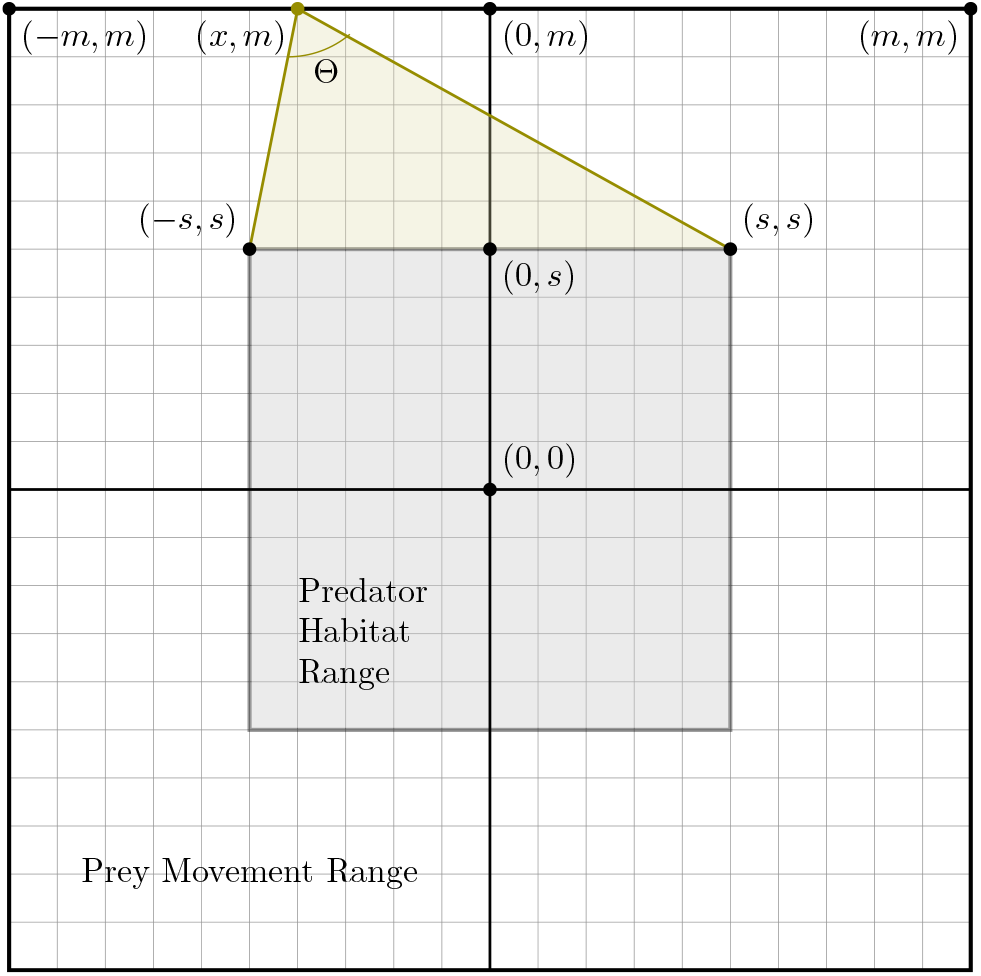
Diagram demonstrating the range of prey headings given their position along a source side that will cross the predator habitat area. In this case, prey originate from the top of the map. A particular prey can have any heading that aims “into” the map; the area spanned by the possible headings that would “hit” the predator habitat for a particular prey at coordinate (*x, m*) is shaded in green.

For a prey position (*x, m*), we want the internal angle Θ of the triangle formed by this point and (−*s, s*) and (*s, s*). We can obtain this by the law of cosines:

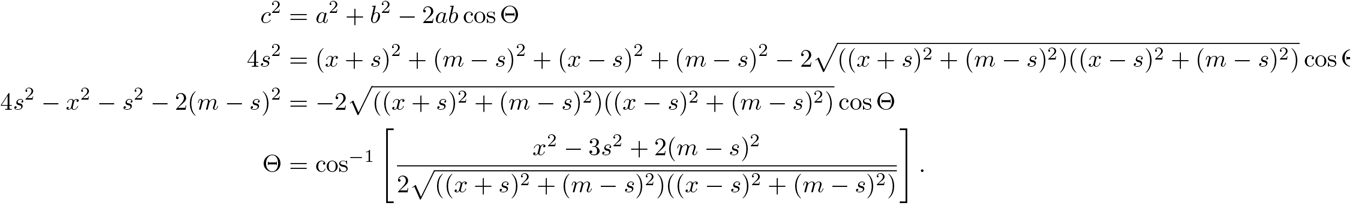

This value of Θ, divided by *π*, is the fraction of possible headings that lead to prey being accessible to predators if a prey enters the map at position *x*. The total fraction of possible headings is obtained by integrating over all *x* between −*m* and *m*.

### B: Relationship between the predator population density and the number of prey per generation

Figure 10 displays several relationships between the predator population density *P* and various quantities related to the number of prey per generation *Y*. Figure 10A demonstrates the motivation for the linear model in eq. 4; the relationship is not really linear, but it is not so nonlinear than another model would do much better. Figure 10B demonstrates that the linear prediction of the number of predators *P*_1_ from eq 4 is indeed close to the observed values of *P*. Again, because this relationship is not perfect, the observations and predictions do not fall on the *P* = *P*_1_ line, but they are close enough for this rough analysis and appear to be scaled by some constant where *P*_1_ is slightly less than *P*. Figure 10C demonstrates that the ratio 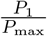 does in fact saturate as *Y* (and therefore *Y* ^*′*^) goes to infinity, as predicted by eq. 5.

**Figure 10:**
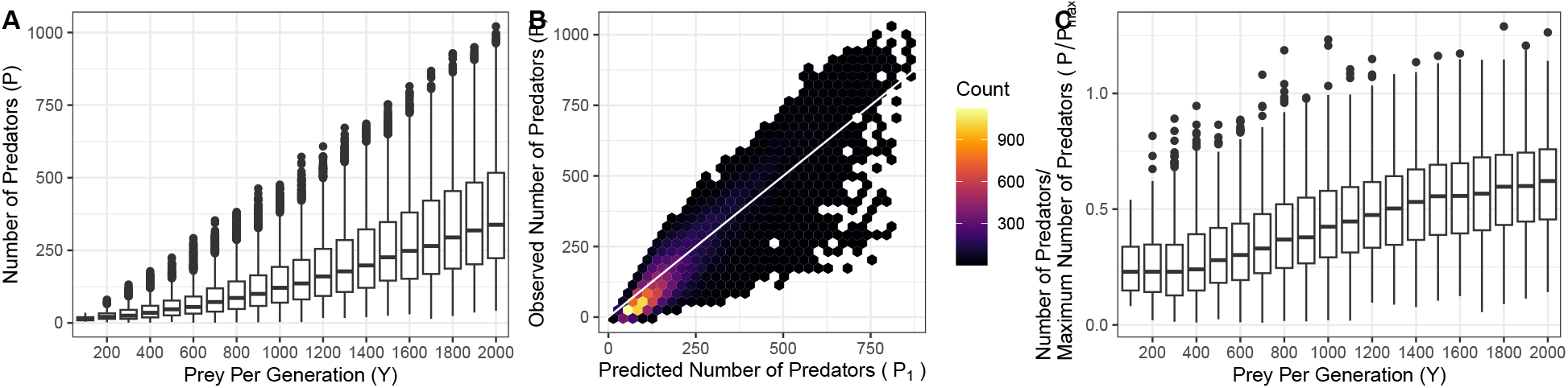
(A) The observed number of predators *P* is approximately linear in the number of prey per generation *Y*, especially for large *Y*. (B) The observed number of predators *P* reasonably closely matches the value predicted from our linear model in eq. 4. (C) The ratio of the number of predators *P* over its theoretical maximum *P*_max_ (eqs. 3 and 5) saturates for high *Y*, which is consistent with our linear model in eq. 4.

